# Multilevel impairment of mitochondrial respiration with sex-specific signatures in inclusion body myositis

**DOI:** 10.64898/2026.02.18.706692

**Authors:** Ibrahim Shammas, Hazem Iaali, Jens O. Watzlawik, Noemi Vidal Folch, Surendra Dasari, Graeme Preston, Thi Kim Oanh Nguyen, Wolfdieter Springer, Tamas Kozicz, Linda Hasadsri, Eugenia Trushina, Ian R. Lanza, Elie Naddaf

## Abstract

**Background:** Oxidative phosphorylation (OXPHOS) is a central function and a key indicator of mitochondrial fitness, yet studies in human tissue remain limited. Inclusion body myositis (IBM) is a progressive myopathy that lies at the intersection of aging, inflammation and mitochondrial dysfunction. We aimed to perform a comprehensive profiling of mitochondrial respiration in muscle tissue from patients with IBM.

**Methods:** A wide battery of complementary tests from RNA level to high-resolution respirometry on permeabilized muscle fibers was performed. The relationship between respiration, mitochondrial content, mitochondrial DNA (mtDNA) abnormalities and mitophagy was examined, along with the correlation with various clinical parameters to determine the clinical significance of the findings.

**Results:** The study included a total of 67 patients with IBM and 45 controls. IBM muscle tissue exhibited reduced maximal respiration per tissue weight in State 3 (high substrates, high ADP) and uncoupled state with decreased coupling efficiency and higher leak control ratios. When adjusting for citrate synthase reflecting mitochondrial content, males had decreased State 3 intrinsic respiration, whereas females had greater intrinsic respiration in leak states. Complex II control ratio strongly correlated with disease duration and severity only in females. IBM was associated with decreased RNA and protein expression of OXPHOS complexes. Complex I activity was decreased mainly in females. IBM samples exhibited lower maximal H_2_O_2_ emission, accompanied by a higher total antioxidant capacity that correlated with disease duration in females. In IBM, there was decreased mtDNA content, and impaired mitophagy, both of which strongly correlated with respirometry measures and markers of disease severity, indicating these pathways are likely interconnected and of clinical significance.

**Conclusion:** IBM is characterized by multilevel impairments in mitochondrial coupling efficiency, revealing several potential therapeutic targets to improve mitochondrial fitness, while accounting for sex-specific differences.

## Introduction

Inclusion body myositis (IBM) is a muscle disease of aging, with no available treatment, characterized by relentlessly progressive muscle weakness, eventually leading to the complete loss of ambulation, marked difficulty swallowing and respiratory insufficiency.^1^ On muscle histopathology, IBM is characterized by the presence of inflammation, accumulation of autophagic vacuoles, protein aggregates, and mitochondrial abnormalities with increased number of cytochrome c oxidase (CCO) negative fibers.^1^ In addition, the accumulation of mitochondrial DNA (mtDNA) deletions has long been recognized in IBM.^2–4^ Nevertheless, the role of mitochondria in IBM pathogenesis is yet to be fully elucidated.

Mitochondrial quality control system consists of a series of interconnected processes, such as proteostasis and mitophagy, that work collectively to maintain mitochondrial fitness and optimize cell survival, through metabolic and respiratory adaptations.^5^ In a recent study, we showed that PINK1- Parkin mitophagy, a protective pathway that is lost in Parkinson’s disease and a key regulator of inflammation, is likely affected in IBM, more severely in males.^6^ Aggregates of phosphorylated ubiquitin (p-S65-Ub), the product of the kinase-ligase pair PINK1-Parkin that serves as the mitophagy marker, occurred mainly in type 2 fibers and the CCO negative fibers were all type 2; type 2 fibers are known to be more prone to mitochondrial and metabolic derangements.^6^ Indeed, muscle samples from patients with IBM demonstrated alterations in central carbon metabolism with sex-specific patterns, sharing similarities with diseases of aging and sarcopenia.^7^ That said, mitochondrial respiration via oxidative phosphorylation remains a key indicator of mitochondrial fitness.^8^ A major limitation in assessing mitochondrial respiration in human samples in general is the limited access to fresh tissue, hence, most published studies were performed either in animal or cellular models.^9^ To an even greater extent, knowledge regarding sex-specific changes in mitochondrial biology including mitochondrial respiration, is limited in humans.^9,10^ To date, no studies have assessed mitochondrial respiration in muscle samples from patients with IBM. Given the access to a large number of muscle biopsies performed at our institution, we aimed to provide a high-resolution profiling of mitochondrial respiration in IBM in a cohort of deeply phenotyped patients, underscoring the clinical significance of the observed changes.

## Methods

### Ethical statement

The Mayo Clinic Institutional Review Board approved the study, which was classified as minimal risk; hence, the need for informed consent was waived. However, in compliance with Minnesota statute 144.335, medical records of patients who had not authorized the use of their information for research were excluded from review. Additionally, the use of residual tissue from muscle biopsy obtained for clinical purposes was approved by the Mayo Clinic Biospecimens Subcommittee.

### Study population

The diagnosis of IBM was established according to the 2013 European Neuromuscular Centre (ENMC) diagnostic criteria for IBM. Controls were patients without myopathy who underwent muscle biopsy to exclude a muscle disorder and had an unremarkable diagnostic workup. All muscle biopsies were performed for clinical purposes. According to our clinical practice standards, an open muscle biopsy was performed in the operating room under conscious sedation after the patient fasted overnight. As high resolution respirometry requires fresh tissue, participants were identified prospectively between 2023 and 2025 (prospective cohort). In addition to the prospectively collected samples, we also included our previously published retrospective cohort of 38 patients with IBM and 22 age- and sex- matched controls with available transcriptomics data in some of the experiments as detailed below.^6,11^

### Clinical parameters

Clinical data were collected through chart review and included patient age at the time of biopsy, sex, disease duration (calculated from symptom onset to the date of biopsy), manual muscle testing (MMT) score and modified ranking scale (MRS). The MMT score represents the total motor strength evaluation of bilateral shoulder abduction, elbow flexion, elbow extension, finger flexion, hip flexion, knee extension, and ankle dorsiflexion. Each muscle is scored from 0 (normal) to 4 (complete paralysis) according to Mayo Clinic scale, and the total score falls between 0 (normal strength) and 56 (complete paralysis) as previously reported.^12^ The MRS, used to evaluate patient’s level of disability, ranges from 0 (no symptoms) to 6 (dead) as follows: MRS 1- symptoms without disability; 2- slight disability with ability to perform baseline activities; 3- moderate disability requiring some help, but able to walk unassisted; 4- moderately severe disability with inability to walk unassisted (wheelchair bound); and 5- severe disability with need for constant nursing care and bedridden.^13^

### Measurement of mitochondrial respiration in permeabilized muscle fibers

Mitochondrial respiration was assessed using the Oxygraph O2K high-resolution fluororespirometer (Oroboros Instruments, Austria) in permeabilized human skeletal muscle fibers **(Figure 1** and **Supplemental Methods)** as previously described, on a prospective cohort identified between 2023 and 2025.^14–16^ Each biopsy sample was run in duplicate: 7-10 mg of tissue per chamber. The muscle tissue was manually dissected under a microscope to achieve high level of fiber separation, then permeabilized by adding saponin. Fibers were added to the Oroboros oxygraph chambers filled with mitochondria respiration media (MiRO5). Oxygen levels were maintained between 200–400 µM throughout the experiment. Various substrates, inhibitors, and uncouplers were sequentially added to assess different mitochondrial respiration states (**Figure 1**): State 2 with high substrates and low ADP, State 3 [Complex I (CI) + Complex II (CII)] with high substrates and high ADP, State 3 (CII) after adding rotenone, State 4 after oligomycin, uncoupled respiration after FCCP, and residual oxygen consumption after Antimycin A. Oxygen consumption (JO_2_) was normalized to the wet tissue weight, and to citrate synthase (CS) activity reflecting intrinsic mitochondrial respiration. Flux control ratios that express respiratory control independent of mitochondrial content or tissue weight, were calculated: respiratory control ratio (RCR) as an indicator of coupling efficiency (State 3: State 4), leak control ratio (State 4: uncoupled respiration) reflecting the proportion of mitochondrial respiration that is attributable to proton leak relative to the maximal electron transport system (ETS) capacity, and CII control ratio (State 3 after rotenone: State 3 after ADP) to assess the relative contribution of CII-supported respiration to the maximal oxidative phosphorylation (OXPHOS) capacity.^14–17^

**Figure 1:**
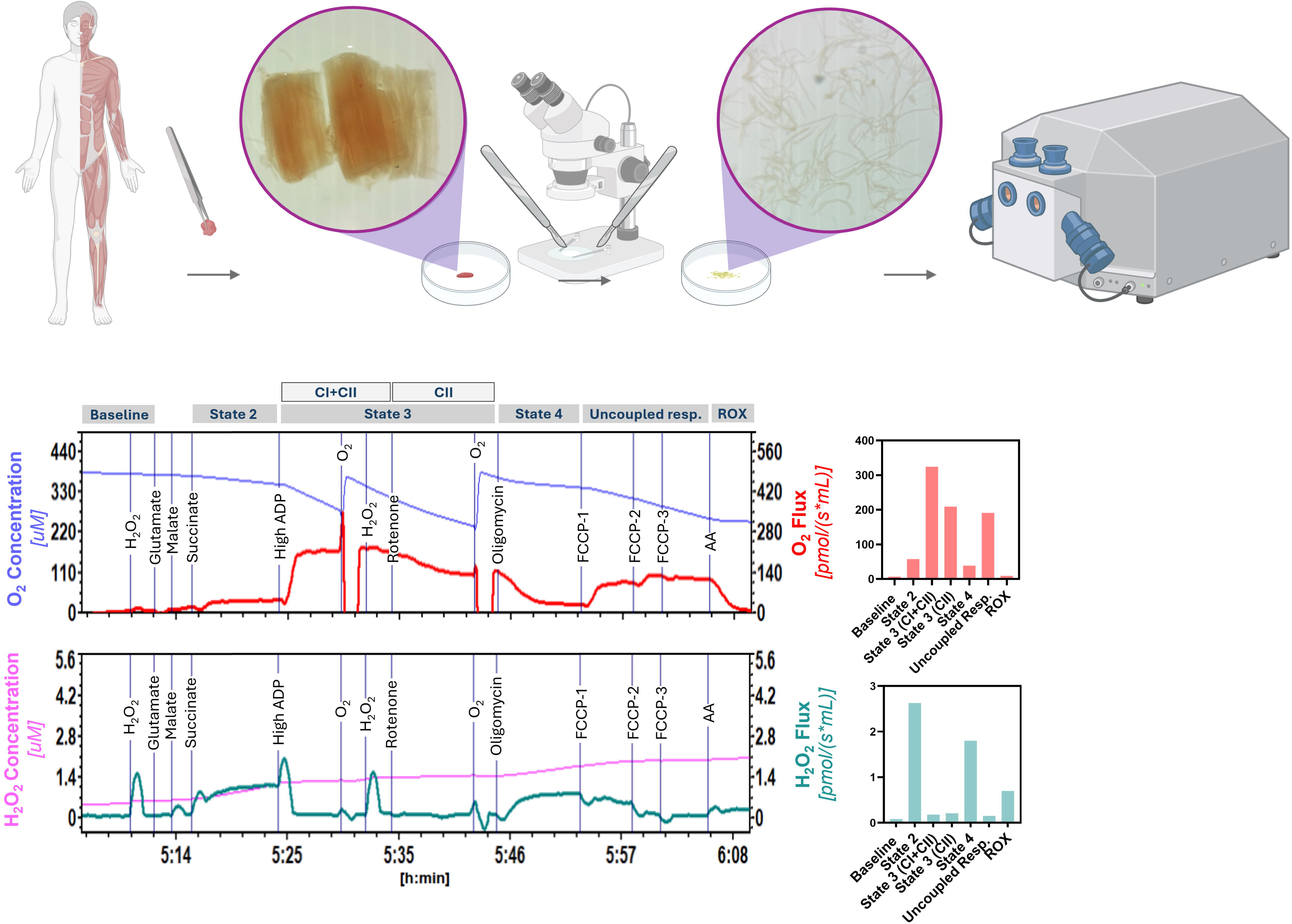
Summary of high-resolution respirometry workflow. Schematic illustration of the experimental workflow for human skeletal muscle processing, including muscle biopsy collection from patients, muscle tissue dissection under a microscope, and representative traces and bar graphs. The upper panel displays real-time oxygen concentration (blue trace) and oxygen flux (red trace), with bar graphs representing raw oxygen flux values in different respiratory states. The lower panel shows real-time hydrogen peroxide (H₂O₂) emission (pink trace) and the corresponding slope (green trace), with representative flux values. *Abbreviations: AA, antimycin A; Resp., respiration; ROX, residual oxygen consumption*.

### Mitochondrial H_2_O_2_ Emission

Hydrogen peroxide (H₂O₂) flux was measured concurrently with respirometry using the Oxygraph O2K equipped with the Fluorescence LED2 module and the H₂O₂-sensitive probe Amplex UltraRed. In the presence of excess superoxide dismutase (SOD), O_2•_^−^ is converted to H_2_O_2_, which reacts with Amplex Red to produce the red fluorescent product, resorufin.^16^ H₂O₂ emission data were normalized to the oxygen consumption of the corresponding respiratory state.^18^ The adopted protocol allows detection of maximal H O emission during State 2.^19^

### RNA transcriptomics

Raw gene-level counts from our recently published whole transcriptomic dataset performed on skeletal muscle samples from the retrospective cohort were used.^6^ The raw gene expression counts were loaded into R statistical programming environment (version 4.1) and normalized using the edgeR software configured to use trimmed means of M-values method.^20^ Normalized expression data was log transformed and submitted to singscore software configured to generate a score quantifying the expression of each electron transport chain complex in each sample using the respective canonical genes.^21^ To evaluate for ROS-related antioxidant defense, we used Reactome detoxification of reactive oxygen species pathway. Processed gene expressions were compared using the default likelihood ratio test (LRT) method. A rank score was computed for each gene in each group comparison using the following formula: - 1*log2(p-value)*sign(fold change). For each group comparison, respective genes and their corresponding rank scores were loaded into Broad’s Gene Set Enrichment Analysis (GSEA) software (version 4.3.3).^22^ The PreRanked method configured with default parameters was utilized to test the association of the gene sets with the differential expression results.

### OXPHOS complexes protein abundance

Measurement of OXPHOS complexes protein levels was performed via western blotting on samples from both cohorts. Equal amounts (15 μg) of proteins per sample were loaded on 4%–20% Criterion TGX™ Precast Protein Gels (Bio-Rad, 4561096) and transferred to a PVDF membrane. Membranes were blocked for 1 h at room temperature in TBS-T with 5% (w/v) non-fat dry milk (NFM), then incubated with the primary antibody overnight at 4 °C. Primary antibodies included Oxidative OXPHOS antibody cocktail (1:500, Abcam, ab110413), which detects NDUFB8 in CI, SDHB in CII, UQCRC2 in CIII, MTCO1 in CIV, and ATP5A in CV, and vinculin (1:500 diluted in TBS-T with 1% NFM, Cell Signaling, 4650). The following secondary antibodies were used: peroxidase-conjugated AffiniPure Goat Anti-Mouse IgG (H+L) (1:5000 diluted in TBS-T with 5% (w/v) Bovine Serum Albumin (BSA), Jackson ImmunoResearch, 115-035-003) and peroxidase-conjugated AffiniPure Goat Anti-Rabbit IgG (H+L) (1:3000 diluted in TBS-T with 5% (w/v) BSA, Jackson ImmunoResearch, 115-035-003). Membranes were developed using the Bio-Rad ChemiDoc Imaging System, and band quantification was carried out with ImageJ software (version 1.54g). Protein expression of each complex was normalized to CII expression in the same lane and to vinculin load. Western blot data were expressed as fold change relative to the corresponding mean value of the controls.

### Electron transport chain complexes activity measurement

The activities of electron transport chain (ETC) complexes were assessed in muscle biopsies homogenate from both patients and controls in the retrospective cohort. Activities of CI, CII, CIII, and CIV were measured using a spectrophotometric assay using a FLUOstar Omega spectrophotometric plate reader (BMG). ^23^ All activities were normalized to CII activity. ^24^ A detailed description of the methods is provided in **Supplemental Methods**.

### Total antioxidant capacity

Total Antioxidant capacity (TAC) was measured in the prospective cohort, using an OxiSelect™ Total Antioxidant Capacity Assay kit as per protocol instructions (Cell Biolabs Inc, San Diego, CA, Cat. NO. STA-360). Briefly, muscle homogenates were incubated with a copper ion reagent. Antioxidants in the homogenates reduce copper ions through a single electron transfer reaction, which subsequently react with a chromogen. Absorbance from the colored complex was measured by spectrophotometry at 490 nm, and TAC was quantified as µM Copper Reducing Equivalents (CRE), normalized to protein content.

### Mitochondrial DNA deletion heteroplasmy levels and mitochondrial DNA content

For mtDNA deletion heteroplasmy levels, a laboratory-developed quantitative digital droplet (ddPCR) assay was performed on the Bio-Rad QX200 Droplet Digital PCR System on the retrospective cohort. The method includes 3 separate ddPCR reactions. In each reaction, DNA extracted from frozen muscle tissue is mixed with primer/probe mixes and ddPCR Supermix.

Mixture is partitioned into nanoliter-sized droplets and targets are amplified by end-point PCR in each droplet, followed by automated measurement of the fluorescence of each droplet in two channels (FAM and HEX). Quantification of mtDNA was performed on both cohorts. QuantaSoft Analysis Pro software allows quantification of multiple targets per reaction and provides copies/20ul well of *RPP30* nuclear gene and *mt-RNR1*, *mt-ND4*, *mt-ND2*, and D-loop mitochondrial genes. For mtDNA content, *mt-RNR1*/*RPP30* ratios were calculated. For mtDNA deletion heteroplasmy levels, various *mt-ND4*/*mt-RNR1*, *mt-ND4*/D-loop genes, *mt-ND4*/*mt-ND2* ratios were used.

### Citrate Synthase Activity

Citrate synthase was measured using previously reported methods on samples from both cohorts, by quantifying the absorbance of thionitrobenzoic acid (TNB) at 412 nm.^25^ TNB is formed during a reaction driven by citrate synthase in the presence of saturating levels of acetyl-CoA, oxaloacetate, and dithionitrobenzoic acid (DNTB). A detailed description of the methods is provided in **Supplementary Methods**.

### Measurement of the mitophagy marker p-S65-Ub

p-S65-Ub levels were measured in muscle lysates from patients with IBM and controls in both cohorts, using a sandwich ELISA via the Meso Scale Discovery (MSD) electrochemiluminescence (ECL) platform, as previously described.^6^

### Statistical Analysis

Continuous data are presented as means with standard deviation (SD) or medians with interquartile range (IQR) as applicable. Categorical variables are presented as proportions and percentages. IBM group was compared to the control group. Statistical analysis was performed using BlueSky statistics v.10.3 software and GraphPadPrism v.10.4. Two-tailed Mann–Whitney tests versus a 2-sample test, and Spearman versus Pearson’s correlation were performed as applicable. Sensitivity analysis adjusting for age difference between patients with IBM and controls was performed via linear regression.

### Data sharing statement

Data not included in this manuscript can be provided by the corresponding author upon reasonable request.

## Results

### Study population

The study population consisted of 67 patients with IBM and 45 controls without a myopathy. This included the prospective cohort with 29 patients with IBM and 23 controls without myopathy, as summarized in **Table 1**. As the prospective cohort included all comers, patients with IBM were approximately two-thirds male and one-third female, a distribution similar to that observed in the general population.^26^ The IBM group was slightly older than the control group. To account for that, sensitivity analysis, adjusting for age, was performed for the key findings and showed relatively similar results as shown in (**Supplemental Table 1**). The retrospective cohort consisted of 38 patients with IBM and 22 age- and sex- matched controls as previously published.^6^

**Table 1:** Patients’ demographics and baseline characteristics for the prospective cohort

| **Variable** | **Inclusion Body Myositis**<br>**(*n* = 29)** | **Controls (*n* = 23)** |
| --- | --- | --- |
| **Age at biopsy (years)** | 64.6 (10.9) | 48.1 (12.1) |
| **Sex** |  |  |
| Females | 11 (38) | 15 (65) |
| Males | 18 (62) | 8 (35) |
| **Race** |  |  |
| American Indian/ Alaskan<br>Native | 0 | 0 |
| Black | 0 | 2 (9) |
| White | 29 (100) | 20 (87) |
| Unknown/not disclosed | 0 | 1 (4) |
| **Disease characteristics** |  |  |
| Disease duration at time<br>of the biopsy (months) | 61 (43.3) | N/A |
| Manual Motor Test score | 17.5 (8) | N/A |
| Modified Rankin Scale | 2 (0.6) | N/A |

Continuous variables are shown as mean (SD), categorical variables as count (percentage).

### Assessment of mitochondrial respiration

High resolution respirometry results are shown in **Figure 2**. There is a decrease in maximal respiration per tissue weight (**Figure 2A)** in IBM in State 3 (CI+II), State 3 (CII) and uncoupled respiration, with no differences in leak states (State 2 and State 4). Similar patterns were seen in males and females **(Supplemental Figure 1A).** When normalizing to CS activity reflecting intrinsic mitochondrial respiration, the differences between IBM and controls become less prominent **(Figure 2B**). However, findings were different by sex where males had lower maximal intrinsic respiration during State 3 (CI+CII and CII) and uncoupled respiration, whereas females had increased intrinsic respiration in leak states (State 2 and State 4) **(Figure 2C).** For flux control ratios, RCR, reflecting coupling efficiency, was decreased, and leak control ratio was increased in IBM compared to controls, with no difference in CII control ratio **(Figure 2D, Supplemental Figure 1B)**.

**Figure 2:**
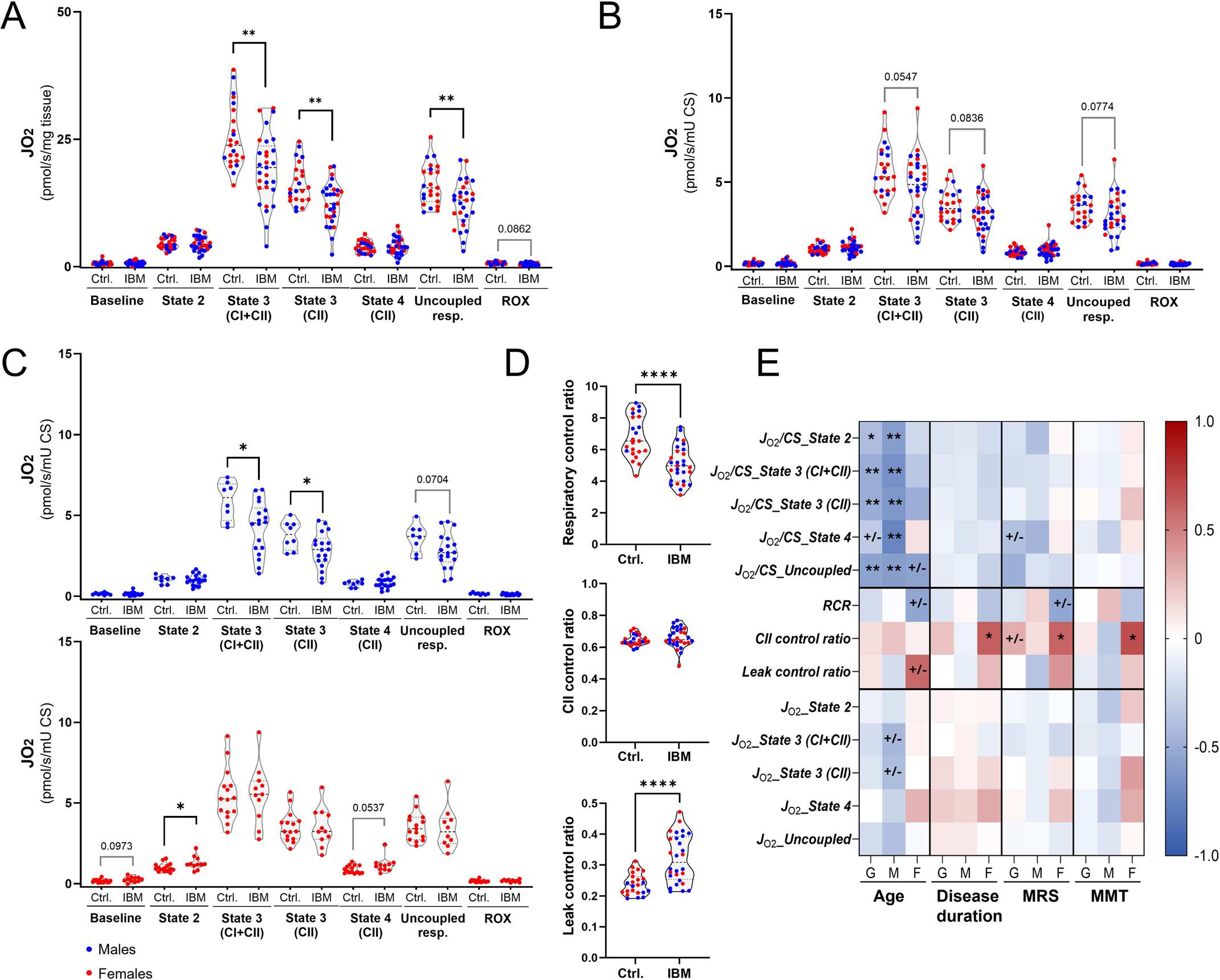
High resolution respirometry in permeabilized muscle fibers. Oxygen consumption rates of permeabilized muscle fibers at each respiratory state in IBM (*n =* 29) and control (*n =* 23) groups normalized by A) tissue weight and B) citrate synthase activity. C) Sex stratification of oxygen consumption rates normalized to citrate synthase activity. D) Flux control ratios. E) Spearman correlations of clinical parameters with flux control ratios and oxygen consumption rates at each respiratory state. *Violin plots show individual data with dashed lines representing medians and interquartile ranges. Unpaired t-tests were used for group comparisons in A, B and D. Mann-Whitney tests were used in C. *P < 0.05, **P < 0.01, ****P < 0.0001, +/-. P values between 0.05 and 0.1. Abbreviations: CI, Complex I; CII, Complex II; CS, Citrate synthase; Ctrl., controls; J_O2_, oxygen consumption rate; IBM, inclusion body myositis; MMT, manual muscle test; MRS, modified Rankin Scale; RCR, respiratory control ratio; ROX, residual oxygen consumption*.

Regarding the clinical correlations of these findings **(Figure 2E)**, age had inverse correlation with all intrinsic respiratory states, most prominently in males. In females, age had inverse correlation with RCR and direct correlation with leak ratio. Disease duration had weak correlations overall except for CII control ratio in females. MMT (muscle weakness severity) and MRS (degree of disability) correlated the most with flux control ratios, most prominently in females. CII ratio in females had one of the strongest correlations with clinical measures, with higher CII respiration correlating with longer disease duration and more severe muscle weakness and disability. Correlations with respiratory ratios often had opposite direction between males and females. Of note, correlations of MMT were the strongest with flux control ratios when stratifying by sex, followed by respiratory per tissue weight with no or very weak correlation with intrinsic respiration.

### RNA expression, protein abundance, and complex activity of OXPHOS complexes

In IBM, OXPHOS complexes pathways were downregulated compared to controls, more significantly in females **(Figure 3** **and Supplemental Figure 2A)**. Likewise, protein levels of all OXPHOS complexes were reduced in IBM, except for CII (**Figure 3)**. CI, CIII and CIV expression was lower in females **(Supplemental Figure 2B)**. Relatively similar results were seen when normalizing to CII expression **(Supplemental Figure 2C, D)**. Relative to CII activity, there is a significant decrease in CI activity in IBM, with no difference in CIII and CIV activities **(Figure 3).** When stratifying by sex, CI/CII activity was significantly decreased in females but not in males **(Supplemental Figure 2F)**.

**Figure 3:**
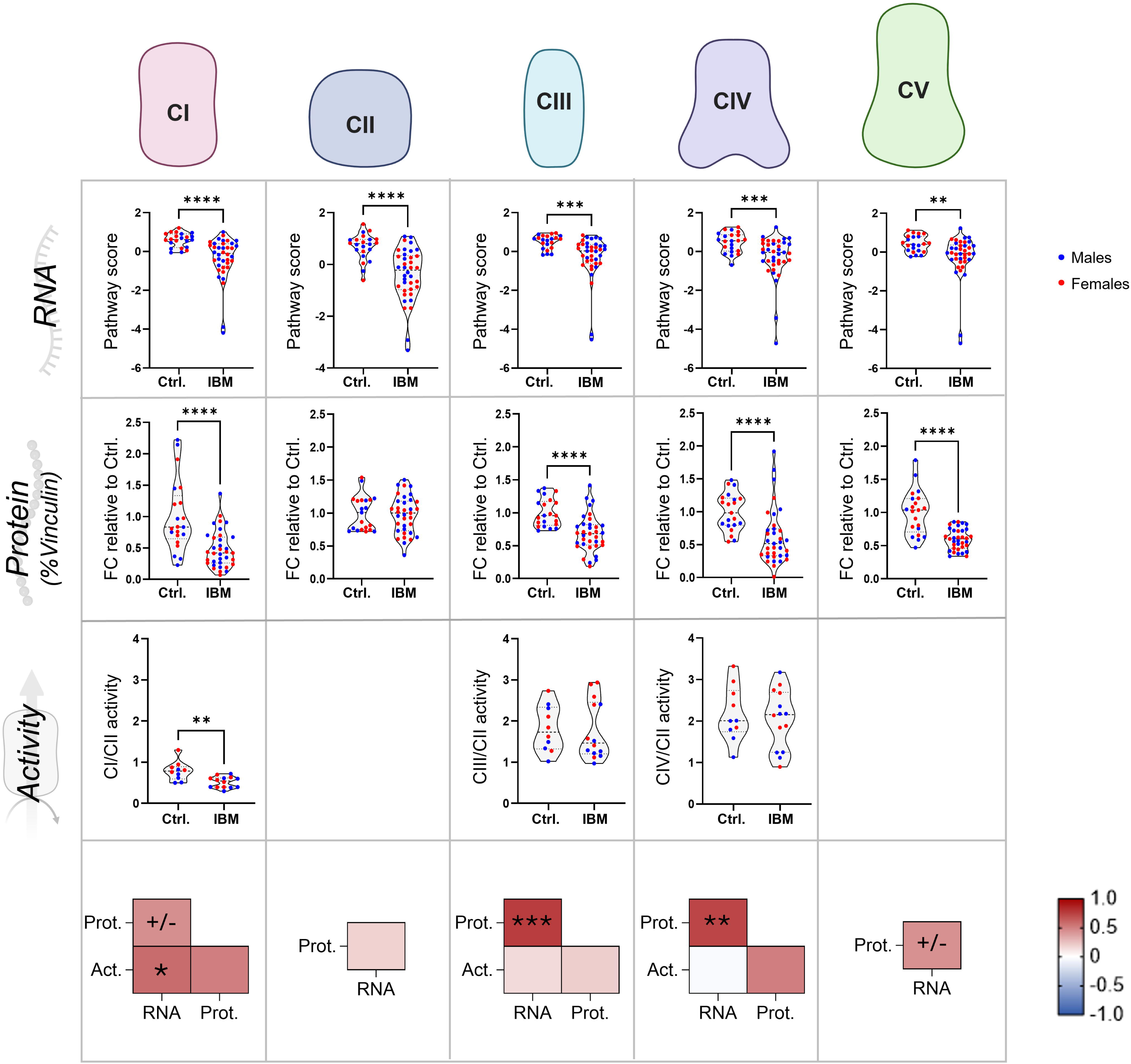
**RNA expression, protein abundance and activity of OXPHOS complexes**. The first row shows RNA pathway scores of OXPHOS complexes I through V in IBM (*n =* 38) and control (*n =* 22) groups from the retrospective cohort. The second row shows the protein abundance of OXPHOS complexes in muscle lysates from patients with IBM (*n =* 35) and controls (*n =* 21) from both cohorts, represented as fold change from the controls’ means. The third row shows the activity of electron transport chain complexes in IBM (*n =* 14) and control (*n =* 10) groups from the retrospective cohort, normalized to Complex II activity. The fourth row shows Spearman correlations for each complex. *Violin plots show individual data with dashed lines representing medians and interquartile ranges. Group comparisons were made using the unpaired t-test in the first row, and Mann Whitney in the second and third. *P < 0.05, **P < 0.01, ***P < 0.001, ***P < 0.0001, +/-. P values between 0.05 and 0.1*. *Abbreviations: Act.: Activity; CI, Complex I; CII, Complex II; CIII, Complex III; CIV, Complex IV; CV, Complex V; CS, citrate synthase; Ctrl., controls; FC, fold change; IBM, inclusion body myositis; Prot., protein*.

There was a strong correlation between RNA expression and protein expression levels in IBM **(Figure 3)**. The sample size for ETC activity was small. Nevertheless, correlation between RNA level, protein level and complex activity was most prominent with CI **(Figure 3)**.

### Assessment of reactive oxygen species production in inclusion body myositis

IBM muscle fibers had lower maximal H_2_O_2_ emission capacity than controls during State 2 (high substrates, low ADP), where maximal H_2_O_2_ emission is typically observed **(Figure 4A** **and Supplemental Figure 3).** Muscle tissue from IBM may have a higher antioxidant capacity given its chronic exposure to oxidative stress. To evaluate for that, we measured the total antioxidant capacity (TAC) in muscle homogenates. The TAC of IBM samples was higher compared to controls **(Figure 4B)**, in both males and females. Furthermore, the detoxification of reactive oxygen species pathway was significantly upregulated at the RNA level, with enrichment of genes related to key antioxidant enzymes and proteins as shown in **Figure 4D**. State 2 H_2_O_2_ emission had a weak correlation with clinical parameters. Conversely, TAC exhibited a strong positive correlation with disease duration, most prominent in females **(Figure 4C)**

**Figure 4:**
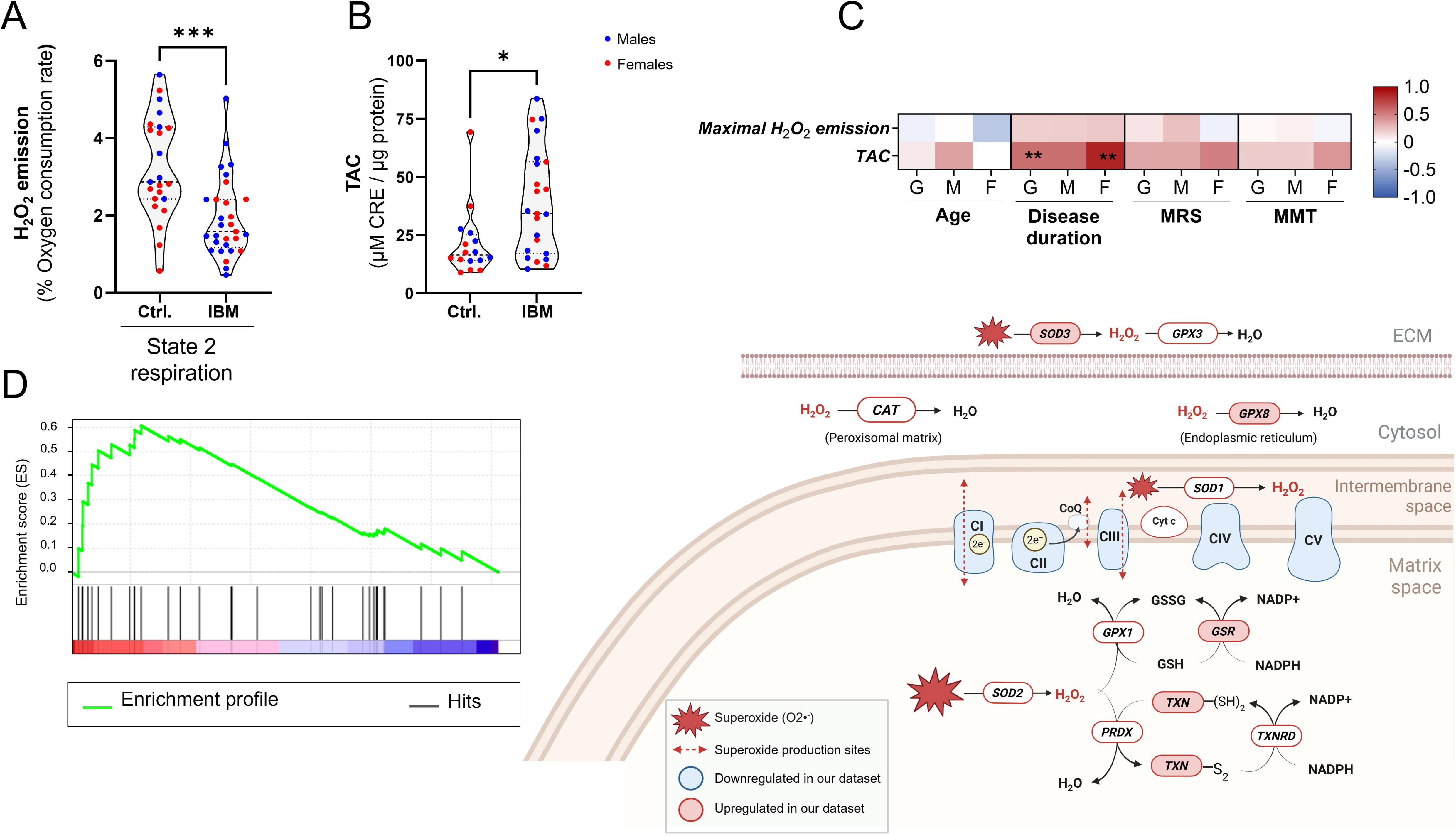
Reactive oxygen species production in IBM. A) Maximal H_2_O_2_ emission in IBM (*n* = 29) and controls (*n* = 23) during State 2 respiration; B) Total antioxidant capacity in muscle homogenates of patients with IBM (*n* = 23) and controls (*n* = 16). C) Spearman correlations of clinical parameters with maximal H_2_O_2_ emission and total antioxidant capacity. D) Left: Gene set enrichment analysis (GSEA) plot of the Reactome “Detoxification of Reactive oxygen species” pathway. Right: Schematic representation of the detoxification of reactive oxygen species and the OXPHOS complexes pathways. *Violin plots show individual data with dashed lines representing medians and interquartile ranges. Group comparisons in (A) and (B) were made using the Mann-Whitney tests. *P < 0.05, **P < 0.01, ***P< 0.001. Abbreviations: CI, Complex I; CII, Complex II; CIII, Complex III; CIV, Complex IV; CV, Complex V; CoQ, coenzyme Q; CRE, copper reducing equivalents; Ctrl., controls; Cyt c, cytochrome c; ECM, extracellular matrix; GSH, reduced glutathione; GSSG, oxidized glutathione; MMT, manual muscle test; MRS, modified Rankin Scale; TAC, total antioxidant capacity*.

### Mitochondrial DNA abnormalities and citrate synthase activity

Muscle samples from IBM group had lower mtDNA content and higher mtDNA deletion heteroplasmy levels compared to controls **(Figure 5A**), which was observed in both males and females **(Supplemental Figure 4A, B)**. In IBM, mtDNA content inversely correlated with MMT, where patients with more severe weakness had lower mtDNA content, most significantly in males. In contrast, mtDNA deletion heteroplasmy levels showed poor correlation with clinical variables **(Figure 5C).**

**Figure 5:**
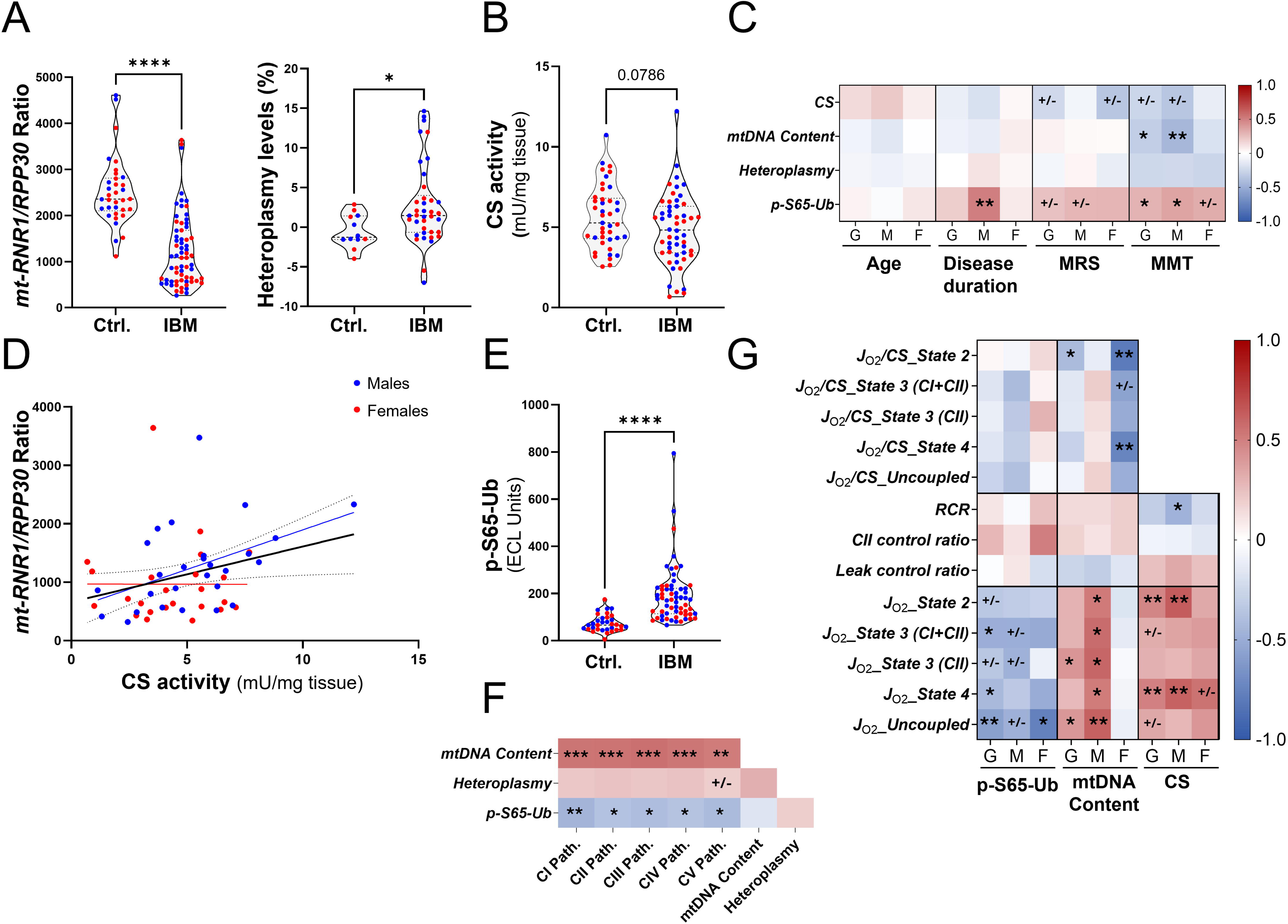
**Mitochondrial DNA abnormalities, and mitophagy defects in inclusion body myositis**. A) Left: Mitochondrial DNA content represented by the *mt-RNR1*/*RPP30* ratio in IBM (*n =* 67) and control (*n =* 34) groups. Right: Mitochondrial DNA deletion heteroplasmy levels in IBM (*n =* 38) and control (*n* =12) groups. B) Tissue mitochondrial content represented by citrate synthase activity, normalized to tissue weight in IBM (*n =* 52) and control (*n =* 41) groups. C) Correlations of clinical parameters with citrate synthase activity, mitochondrial DNA content, mitochondrial DNA deletion heteroplasmy level and p-S65-Ub levels. D) Correlation between tissue citrate synthase activity and mitochondrial DNA content (*mt-RNR1*/*RPP30 ratio).* Fitted lines are shown as black for the whole group (black) with dotted 95% confidence interval, and stratified by sex (blue for male, red for female). E) p-S65-Ub levels in IBM (*n =* 62) and controls (*n =* 33). F) Spearman correlations between complex RNA pathway scores, mitochondrial DNA content, mtDNA deletion heteroplasmy levels and p-S65-Ub levels. G) Spearman correlations of oxygen consumption rates and flux control ratios with p-S65-Ub levels, mitochondrial DNA content and citrate synthase activity. *Violin plots show individual data with dashed lines representing medians and interquartile ranges. Group comparisons were made using unpaired t-tests in A, B, E, with Welch’s correction in E. mtDNA deletion heteroplasmy levels group comparisons were done using the Mann-Whitney test. *P < 0.05, **P < 0.01, ***P < 0.001, ***P < 0.0001; +/-. P values between 0.05 and 0.1. Abbreviations: CI, Complex I; CII, Complex II; CS, citrate synthase activity; Ctrl., controls; JO2, oxygen consumption rate; MMT, manual muscle test; MRS, modified Rankin Scale; mtDNA, mitochondrial DNA; path., pathway score; RCR, respiratory control ratio*.

The CS activity per tissue weight was mildly decreased in IBM **(Figure 5B)**, more notably in females **(Supplemental Figure 4C);** the difference between groups was statistically significant after adjusting for age **(Supplemental Table 1).** Clinically, CS activity had moderate inverse correlations with MRS and MMT **(Figure 5C).**

Both citrate synthase activity and mtDNA content may be used to reflect mitochondrial content in skeletal muscle. There was a significant correlation between mtDNA content and CS activity (slope 93.82, *p*=0.037) in IBM **(Figure 5D)**, mostly in males. However, a floor effect was observed with mtDNA content with multiple samples clustering at low values despite having various levels of CS activity. Furthermore, mtDNA content had a strong correlation with RNA expression of all complexes **(Figure 5F)**.

### Relationship between mitochondrial respiration, mtDNA abnormalities, and mitophagy defects in inclusion body myositis

As p-S65-Ub is a PINK1-PRKN mitophagy marker that reflects the burden of accumulated damaged mitochondria in the cell, we sought to evaluate the correlation of p-S65-Ub with mitochondrial respiration.^27,28^ P-S65-Ub levels were elevated in samples from patients with IBM, more notably in males **(Figure 5E** **and Supplemental Figure 4D).** p-S65-Ub levels correlated with clinical parameters, most significantly in males **(Figure 5C)**. Furthermore, p-S65-Ub levels showed strong correlation with the RNA expression of all complexes **(Figure 5F)**.

Regarding the correlations with respirometry parameters in IBM, p-S65-Ub had the most significant correlations with respiration per tissue weight, especially with uncoupled respiration, where samples with higher p-S65-Ub levels (greater mitophagy dysregulation) had lower respiration per tissue weight and lower ETS capacity **(Figure 5G).** Among flux control ratios, CII respiratory ratio in females showed the strongest correlation with p-S65-Ub levels in IBM, although it did not reach statistical significance. Regarding mtDNA content, there were striking sex specific differences. In males, maximal respiration per tissue weight strongly correlated with mtDNA content in all states. In contrast, females had inverse correlations between mtDNA content and intrinsic respiration during leak states, indicating females with lower mtDNA content had higher intrinsic respiration during leak states **(Figure 5G).** Lastly, CS activity correlated with respiration per tissue weight, especially in leak states, and with RCR in males.

## Discussion

In this study, we showed altered mitochondrial respiration in IBM, characterized by decreased maximal State 3 respiration and uncoupled respiration per tissue weight, as well as impaired coupling efficiency and leak control, in both males and females. Furthermore, males showed decreased maximal State 3 intrinsic mitochondrial respiration, whereas females had higher intrinsic respiration during leak states. Age significantly correlated with intrinsic mitochondrial respiration in IBM, mainly in males. Among disease severity measures (MRS and MMT), flux control ratios (RCR and Leak control ratio) had the strongest correlations, often with opposite direction of association between males and females. Previous studies using fibroblasts derived from a small number of patients with IBM have shown lower oxygen consumption rate in IBM compared to controls.^29,30^ However, studies evaluating mitochondrial respiration in muscle disorders are generally rare. One study reported lower maximal respiration in muscle tissue from dermatomyositis patients using a Clark electrode, and another study showed lower maximal respiration but with normal coupling efficiency (State 3/State 2 with glutamate and malate) in myotonic dystrophy type 1 (DM1) using Oroboros oxygraph.^31,32^ Likewise, sex differences in mitochondrial biology in humans, including respiration, are poorly explored as summarized in a recent meta-analysis, with only 22 studies identified in the context of diabetes, aging and exercise physiology, showing no clear differences in maximal respiration between males and females.^9,10^ In IBM, there was decreased RNA and protein expression of mitochondrial respiration complexes, more notable in females. Despite the small sample size for ETC activity measurement, there was a significant decrease in CI activity relative to CII, most significantly in females. This is in line with respirometry results, where females, but not males, with longer disease duration and more severe muscle weakness exhibited more respiration through CII. It is possible that the decreased respiration through CI and increased respiration through leak states in females represent a compensatory mechanism to cope with elevated oxidative stress, as higher proton leak can help reduce ROS generation, or to improve mitochondrial fitness via other mechanisms.^33^

IBM muscle fibers had lower maximal ROS-producing capacity per O_2_ consumption than controls. Studies about ROS production in human muscle are limited with a recent study reporting lower maximal ROS production capacity in DM1.^32^ One potential explanation is that muscle fibers in IBM, chronically exposed to oxidative stress, develop better defense mechanisms. This was shown on RNA level with upregulation of the detoxification of ROS pathway and the higher total antioxidant capacity in IBM in our study. This finding is consistent with TAC measurement in IBM fibroblasts.^29,34^ An increase in DJ-1, another antioxidant protein implicated in Parkinson’s disease, has been reported in IBM muscle samples.^35^ Furthermore, a strong positive correlation was observed between TAC and disease duration in our data, particularly among females. We have previously shown that most of the upregulated female sex-specific genes, on a whole transcriptomic level in IBM muscle samples, are related to stress response.^6^

Mitochondrial DNA abnormalities, characterized by increased mtDNA deletions and reduced mtDNA content, have been described in IBM and confirmed in our study.^36,37^ We built on this to determine the relationship between mtDNA abnormalities, clinical characteristics, mitochondrial content and mitochondrial respiration in this study. First, mtDNA content showed strong inverse correlation with the severity of muscle weakness, especially in males. mtDNA content also correlated with CS activity. Determining the best marker for mitochondrial content in IBM is out of the scope of this study. However, there appears to be a floor effect with mtDNA content with multiple samples clustering at low values despite having various levels of CS activity. This is an interesting finding and may be suggestive of an increased number of mitochondria that have released their mtDNA. The release of mtDNA is a main trigger for inflammasome activation.^6^ Furthermore, a recent study demonstrated elevated levels of circulating cell-free mtDNA in IBM.^38^ Mitophagy, is a key element of mitochondrial quality control, which ensures the recycling of damaged mitochondria that would otherwise trigger inflammatory pathways, such as inflammasomes.^39^ First, we confirmed that mitophagy is altered in IBM in a larger cohort with significant correlations of the mitophagy label p-S65-Ub with disease severity and duration, particularly in males. Next, we demonstrated that mitophagy dysregulation and impaired mitochondrial respiration in muscle tissue are interconnected, as evidenced by the significant correlations with respirometry parameters, particularly in males. Similarly, mtDNA content is strongly linked to mitochondrial respiration in a sex-specific pattern with strong association with maximal respiration per tissue in all states in males, and intrinsic respiration in leak states in females; a recurrent finding observed across the study. Furthermore, there was a consistent pattern suggestive of females having lower mitochondrial content (CS activity, mtDNA content, RNA expression of ETC-related genes). Taken together, females with IBM tend to have relatively lower mitochondrial content, less affected mitophagy, and more preserved maximal intrinsic respiration but with higher leak, compared to males. Studies in animal and cellular models showed a link between altered mitophagy and defective mitochondrial respiration, with interventions aimed to enhance mitophagy subsequently improving mitochondrial respiration in cardiac and skeletal muscle.^40–44^ Beyond neurodegenerative diseases, enhancing mitophagy could be a potential therapeutic intervention to also be explored in IBM in future studies.^45^

Limitations of the study include the retrospective nature of the chart review, limiting the clinical parameters that can be abstracted. The observed correlations cannot infer causal relationships, and the performance of more in-depth mechanistic studies is hindered by the lack of a disease model for IBM.^1^ Nevertheless, performing all the experiments on patient-derived samples and determining the clinical significance of the findings would enhance the translational potential of this study. Lastly, despite the relatively larger sample size given the disease rarity, powering for subgroup analysis by sex or multiple comparisons from a statistical standpoint would markedly limit study feasibility. All comparisons beyond IBM versus control groups were considered exploratory and *p*-values between 0.05 and 0.1 were still highlighted for the reader. Given the limited amount of tissue available, not all experiments were performed on all samples; however, sample sizes for each experiment are mentioned throughout the results section.

## Conclusion

Decreased coupling efficiency in IBM is indicative of compromised mitochondrial fitness and is likely due to the cumulative effect of multiple factors, some of which were addressed in this study, providing multiple potentials points for intervention. Subsequently, future studies aiming to enhance mitochondrial fitness in IBM should be considered, taking into account sex-specific differences.

## Author contributions

Study concept, and supervision: EN. Study design: E.N., E.T., and I.R. All experiments were performed by I.S., H.I, and E.N, except for complex activity measurements by G.P. and T.K., mitochondrial DNA abnormalities by N.V.F, and p-S65-Ub measurements by J.O.W. and W.S.

T.K.O.N. helped with western blotting. Bioinformatical and statistical analysis: S. D. and E.N. Manuscript drafting: I.S., H.I., and E.N. Data interpretation, manuscript revision for intellectual content and approval of final manuscript: all authors.

## References

1. Lilleker JB, Naddaf E, Saris CGJ, Schmidt J, de Visser M, Weihl CC. 272nd ENMC international workshop: 10 Years of progress - revision of the ENMC 2013 diagnostic criteria for inclusion body myositis and clinical trial readiness. 16-18 June 2023, Hoofddorp, The Netherlands. Neuromuscul Disord. Apr 2024;37:36–51. doi:10.1016/j.nmd.2024.03.001

2. Rygiel KA, Miller J, Grady JP, Rocha MC, Taylor RW, Turnbull DM. Mitochondrial and inflammatory changes in sporadic inclusion body myositis. Neuropathol Appl Neurobiol. Apr 2015;41(3):288–303. doi:10.1111/nan.12149

3. Hedberg-Oldfors C, Lindgren U, Basu S, et al. Mitochondrial DNA variants in inclusion body myositis characterized by deep sequencing. *Brain pathology (Zurich*, Switzerland). May 2021;31(3):e12931. doi:10.1111/bpa.12931

4. Dahlbom K, Lindberg C, Oldfors A. Inclusion body myositis: morphological clues to correct diagnosis. Neuromuscul Disord. Nov 2002;12(9):853–7. doi:10.1016/s0960-8966(02)00098-6

5. Bennett CF, Latorre-Muro P, Puigserver P. Mechanisms of mitochondrial respiratory adaptation. Nature Reviews Molecular Cell Biology. 2022/12/01 2022;23(12):817-835. doi:10.1038/s41580-022-00506-6

6. Naddaf E, Nguyen TKO, Watzlawik JO, et al. NLRP3 Inflammasome Activation and Altered Mitophagy Are Key Pathways in Inclusion Body Myositis. J Cachexia Sarcopenia Muscle. Feb 2025;16(1):e13672. doi:10.1002/jcsm.13672

7. Naddaf E, Shammas I, Dasari S, et al. Mitochondria-centred metabolomic map of inclusion body myositis: sex-specific alterations in central carbon metabolism. Annals of the rheumatic diseases. 2025/05/30/ 2025;10.1016/j.ard.2025.05.003

8. Smolina N, Khudiakov A, Kostareva A. Assaying Mitochondrial Respiration as an Indicator of Cellular Metabolism and Fitness. Methods Mol Biol. 2023;2644:3–14. doi:10.1007/978-1-0716-3052-5_1

9. Junker A, Wang J, Gouspillou G, et al. Human studies of mitochondrial biology demonstrate an overall lack of binary sex differences: A multivariate meta-analysis. Faseb j. Feb 2022;36(2):e22146. doi:10.1096/fj.202101628R

10. Ferguson EJ, Pacitti LJ, Bureau J, et al. Biological sex does not impact intrinsic mitochondrial respiration supported by complexes I and II in human skeletal muscle. Exp Physiol. Sep 2025;110(9):1302–1315. doi:10.1113/ep092551

11. Naddaf E, Shammas I, Dasari S, et al. Mitochondria-centred metabolomic map of inclusion body myositis: sex-specific alterations in central carbon metabolism. Ann Rheum Dis. Aug 2025;84(8):1375–1386. doi:10.1016/j.ard.2025.05.003

12. Pinto MV, Laughlin RS, Klein CJ, Mandrekar J, Naddaf E. Inclusion body myositis: correlation of clinical outcomes with histopathology, electromyography and laboratory findings. *Rheumatology (Oxford*, England). May 30 2022;61(6):2504–2511. doi:10.1093/rheumatology/keab754

13. van Swieten JC, Koudstaal PJ, Visser MC, Schouten HJ, van Gijn J. Interobserver agreement for the assessment of handicap in stroke patients. Stroke. May 1988;19(5):604–7. doi:10.1161/01.str.19.5.604

14. Lanza IR, Zabielski P, Klaus KA, et al. Chronic caloric restriction preserves mitochondrial function in senescence without increasing mitochondrial biogenesis. Cell metabolism. Dec 5 2012;16(6):777–88. doi:10.1016/j.cmet.2012.11.003

15. Gnaiger E. Capacity of oxidative phosphorylation in human skeletal muscle: new perspectives of mitochondrial physiology. Int J Biochem Cell Biol. Oct 2009;41(10):1837–45. doi:10.1016/j.biocel.2009.03.013

16. Anderson EJ, Lustig ME, Boyle KE, et al. Mitochondrial H2O2 emission and cellular redox state link excess fat intake to insulin resistance in both rodents and humans. J Clin Invest. Mar 2009;119(3):573–81. doi:10.1172/JCI37048

17. Pesta D, Gnaiger E. High-resolution respirometry: OXPHOS protocols for human cells and permeabilized fibers from small biopsies of human muscle. Methods Mol Biol. 2012;810:25–58. doi:10.1007/978-1-61779-382-0_3

18. Abid H, Ryan ZC, Delmotte P, Sieck GC, Lanza IR. Extramyocellular interleukin-6 influences skeletal muscle mitochondrial physiology through canonical JAK/STAT signaling pathways. Faseb j. Nov 2020;34(11):14458–14472. doi:10.1096/fj.202000965RR

19. Nolfi-Donegan D, Braganza A, Shiva S. Mitochondrial electron transport chain: Oxidative phosphorylation, oxidant production, and methods of measurement. Redox biology. Oct 2020;37:101674. doi:10.1016/j.redox.2020.101674

20. Robinson MD, McCarthy DJ, Smyth GK. edgeR: a Bioconductor package for differential expression analysis of digital gene expression data. *Bioinformatics (Oxford*, England). Jan 1 2010;26(1):139–40. doi:10.1093/bioinformatics/btp616

21. Foroutan M, Bhuva DD, Lyu R, Horan K, Cursons J, Davis MJ. Single sample scoring of molecular phenotypes. BMC bioinformatics. Nov 6 2018;19(1):404. doi:10.1186/s12859-018-2435-4

22. Subramanian A, Tamayo P, Mootha VK, et al. Gene set enrichment analysis: a knowledge-based approach for interpreting genome-wide expression profiles. Proc Natl Acad Sci U S A. Oct 25 2005;102(43):15545–50. doi:10.1073/pnas.0506580102

23. Rodenburg RJ, Schoonderwoerd GC, Tiranti V, et al. A multi-center comparison of diagnostic methods for the biochemical evaluation of suspected mitochondrial disorders. Mitochondrion. Jan 2013;13(1):36–43. doi:10.1016/j.mito.2012.11.004

24. Frazier AE, Vincent AE, Turnbull DM, Thorburn DR, Taylor RW. Assessment of mitochondrial respiratory chain enzymes in cells and tissues. Methods Cell Biol. 2020;155:121–156. doi:10.1016/bs.mcb.2019.11.007

25. Wang Y, Walsh SW. Placental mitochondria as a source of oxidative stress in pre-eclampsia. Placenta. Nov 1998;19(8):581–6. doi:10.1016/s0143-4004(98)90018-2

26. Naddaf E, Shelly S, Mandrekar J, et al. Survival and associated comorbidities in inclusion body myositis. *Rheumatology (Oxford*, England). May 5 2022;61(5):2016–2024. doi:10.1093/rheumatology/keab716

27. Narendra DP, Youle RJ. The role of PINK1-Parkin in mitochondrial quality control. Nat Cell Biol. Oct 2024;26(10):1639–1651. doi:10.1038/s41556-024-01513-9

28. Fiesel FC, Ando M, Hudec R, et al. (Patho-)physiological relevance of PINK1-dependent ubiquitin phosphorylation. EMBO Rep. Sep 2015;16(9):1114–30. doi:10.15252/embr.201540514

29. Catalan-Garcia M, Garcia-Garcia FJ, Moreno-Lozano PJ, et al. Mitochondrial Dysfunction: A Common Hallmark Underlying Comorbidity between sIBM and Other Degenerative and Age-Related Diseases. J Clin Med. May 13 2020;9(5)doi:10.3390/jcm9051446

30. Oikawa Y, Izumi R, Koide M, et al. Mitochondrial dysfunction underlying sporadic inclusion body myositis is ameliorated by the mitochondrial homing drug MA-5. PloS one. 2020;15(12):e0231064. doi:10.1371/journal.pone.0231064

31. Meyer A, Laverny G, Allenbach Y, et al. IFN-beta-induced reactive oxygen species and mitochondrial damage contribute to muscle impairment and inflammation maintenance in dermatomyositis. Acta Neuropathol. Oct 2017;134(4):655–666. doi:10.1007/s00401-017-1731-9

32. Marcangeli V, Girard-Côté L, Di Leo V, et al. A 12-Week Strength Training Improves Mitochondrial Respiration, H(2)O(2) Emission and Skeletal Muscle Integrity in Women With Myotonic Dystrophy Type 1. Acta Physiol (Oxf). Dec 2025;241(12):e70135. doi:10.1111/apha.70135

33. Brookes PS. Mitochondrial H(+) leak and ROS generation: an odd couple. Free Radic Biol Med. Jan 1 2005;38(1):12–23. doi:10.1016/j.freeradbiomed.2004.10.016

34. Cantó-Santos J, Valls-Roca L, Tobías E, et al. Unravelling inclusion body myositis using a patient-derived fibroblast model. J Cachexia Sarcopenia Muscle. Apr 2023;14(2):964–977. doi:10.1002/jcsm.13178

35. Terracciano C, Nogalska A, Engel WK, Wojcik S, Askanas V. In inclusion-body myositis muscle fibers Parkinson-associated DJ-1 is increased and oxidized. Free Radic Biol Med. Sep 15 2008;45(6):773–9. doi:10.1016/j.freeradbiomed.2008.05.030

36. Lindgren U, Hedberg-Oldfors C, Pullerits R, Lindberg C, Oldfors A. Inclusion body myositis with early onset: a population-based study. J Neurol. Jul 27 2023;doi:10.1007/s00415-023-11878-w

37. Catalán-García M, Garrabou G, Morén C, et al. Mitochondrial DNA disturbances and deregulated expression of oxidative phosphorylation and mitochondrial fusion proteins in sporadic inclusion body myositis. Clinical Science. 2016;130(19):1741–1751. doi:10.1042/cs20160080

38. Kleefeld F, Cross E, Lagos D, et al. Mitochondrial damage is associated with an early immune response in inclusion body myositis. Brain. Sep 3 2025;148(9):3199–3214. doi:10.1093/brain/awaf118

39. Marchi S, Guilbaud E, Tait SWG, Yamazaki T, Galluzzi L. Mitochondrial control of inflammation. Nat Rev Immunol. Mar 2023;23(3):159–173. doi:10.1038/s41577-022-00760-x

40. Siddall HK, Yellon DM, Ong SB, et al. Loss of PINK1 increases the heart’s vulnerability to ischemia-reperfusion injury. PloS one. 2013;8(4):e62400. doi:10.1371/journal.pone.0062400

41. Luan P, D’Amico D, Andreux PA, et al. Urolithin A improves muscle function by inducing mitophagy in muscular dystrophy. Sci Transl Med. Apr 7 2021;13(588)doi:10.1126/scitranslmed.abb0319

42. Wang SH, Zhu XL, Wang F, et al. LncRNA H19 governs mitophagy and restores mitochondrial respiration in the heart through Pink1/Parkin signaling during obesity. Cell Death Dis. May 28 2021;12(6):557. doi:10.1038/s41419-021-03821-6

43. Gouspillou G, Godin R, Piquereau J, et al. Protective role of Parkin in skeletal muscle contractile and mitochondrial function. J Physiol. Jul 2018;596(13):2565–2579. doi:10.1113/jp275604

44. Picca A, Faitg J, Auwerx J, Ferrucci L, D’Amico D. Mitophagy in human health, ageing and disease. Nat Metab. Dec 2023;5(12):2047–2061. doi:10.1038/s42255-023-00930-8

45. Antico O, Thompson PW, Hertz NT, Muqit MMK, Parton LE. Targeting mitophagy in neurodegenerative diseases. Nature reviews Drug discovery. Apr 2025;24(4):276–299. doi:10.1038/s41573-024-01105-0

