## Supplemental Methods and Figures for "Multilevel impairment of mitochondrial respiration with sex-specific signatures in inclusion body myositis"

**High Resolution Respirometry**

The Oroboros O2K high-resolution fluoro-respirometer was used to assess mitochondrial respiration in permeabilized muscle fibers from human skeletal muscles biopsies, as previously reported [1-3]. Right after collection, the muscle sample was put on ice in BIOPS buffer, containing 50 mM K + -MES, 20 mM taurine, 0.5 mM dithiothreitol, 6.56 mM MgCl 2 , 5.77 mM ATP, 15 mM phosphocreatine, 20 mM imidazole, pH 7.1, adjusted with 5 N KOH at 0 °C, 10 mM Ca–EGTA buffer. Each sample was run in duplicate, one piece (7-12 mg) for each chamber. Each piece is placed on ice on a Petri dish with 2 mL of BIOPS buffer. Muscle fibers were dissected using two pairs of scalpels under a dissection microscopy, to reach a high level of fibers segregation. Afterwards, 20 uL of Saponin (5mg/mL), a plasma membrane permeabilizer, were added to each Petri dish while shaking for 30 minutes. Meanwhile, Oroboros chamber, was filled with 2.5 mL of respiration media (MiRO5), containing 0.5 mM EGTA, 3mM MgCl2*6H2O, 60 mM potassium lactobionate, 20 mM taurine, 10 mM KH2PO4, 20 mM HEPES, 110 mM Sucrose, and 1 g/l fatty acid free BSA. (MiR05; Oroboros, Innsbruck, Austria). The chambers were initially calibrated with ambient air. Blebbistatin, a fiber contraction inhibitor, was added, followed by Amplex Red (AMR), horseradish peroxidase (HRP), superoxide dismutase (SOD) and stepwise H₂O₂ calibration. Oxygen levels were maintained between 200- 400μM throughout all respiration measurements. Permeabilized fibers were added to each chamber. This was followed by serial additions of various substrates, inhibitors and uncouplers to investigate various states of mitochondrial respiration. The experiment began with the addition of glutamate, malate, and succinate to evaluate State 2, followed by high ADP to evaluate State 3 (CI+II), rotenone (complex I inhibitor) to evaluate State 3 (CII), oligomycin (Complex V inhibitor) to evaluate State 4, then stepwise titration of carbonylcyanide-4-(trifluoromethoxy)-phenylhydrazone (FCCP) (uncoupler) to evaluate uncoupled respiration, and finally, antimycin A (complex III inhibitor) to evaluate non-mitochondrial respiration.

**Electron transport chain activity measurement**

Frozen muscle samples were homogenized at 5% (20mg/400uL) in 10 mM Tris-HCl (pH 7.6), 2mM EDTA, 25mM Sucrose, and 50U/mL Heparin using a bead mill homogenizer and 1.5 mL microtubes pre-filled with 1.4mm ceramic beads (Omni International). The activities of mitochondrial complexes I (CI), II (CII), III (CIII), and IV (CIV) were assessed using a spectrophotometric enzyme activity assay performed on a FLUOstar Omega spectrophotometric plate reader (BMG). Complex activities were expressed relative to CII.

- Complex I (CI): Tissue homogenates were incubated in a potassium phosphate (K_2_HPO_4_) buffered solution of bovine serum albumin (BSA) in the presence of nicotinamide adenine dinucleotide (NADH), ubiquinone (CoQ), and the artificial electron receptor 2,6-dichlorophenolindophenol (DCPIP). Ubiquinone reduced to ubiquinol (QH_2_) by the oxidation of NADH to NAD^+^ by CI rapidly reduces DCPIP (blue) to DCIPIH_2_ (colorless). The NADH oxidation was assayed through spectrophotometric measurement of the extinction of DCPIP absorption at 600nm over 17 minutes. Non-specific NADH oxidation was determined by simultaneously assaying each tissue homogenate in the presence of the potent CI inhibitor rotenone, and CI activity was calculated by subtracting the non-specific (rotenone-inhibited) NADH oxidation from the total (rotenone-uninhibited) NADH oxidation.
- Complex II (CII): Tissue homogenates were incubated in a K_2_HPO_4_ buffered solution of BSA, EDTA, and sodium azide (NaAz) in the presence of succinate, decylubiquinone (DUB), adenosine triphosphate (ATP) and DCPIP. DUB reduced to QH2 by the oxidation of succinate by CII rapidly oxidizes DCPIP (blue) to DCIPIH_2_ (colorless). Succinate oxidation was assayed through spectrophotometric measurement of the extinction of DCPIP absorbance at 600nm for 15 minutes. Non-specific reduction of DCPIP was corrected for by simultaneously assaying each tissue homogenate in the presence of the potent CII inhibitor malonate, and CII activity was calculated by subtracting the non-specific (malonate-inhibited) succinate oxidation from the total (malonate-uninhibited) succinate oxidation.
- Complex III (CIII): Tissue homogenates were incubated in a K_2_HPO_4_ buffered solution of EDTA, sodium azide (NaAz), and polysorbate 20 in the presence of reduced DUB (DUH_2_) and cytochrome C (CytC). Reduction of CytC by CIII via oxidation of DUH_2_ was assayed via spectrophotometric measurement of reduced CytC at 550nm over 15 minutes. Non-specific reduction of CytC was corrected for by first assaying for CytC reduction in the presence of DUH2 but the absence of the homogenate.
- Complex IV (CIV): Tissue homogenates were incubated in a K_2_HPO_4_ buffered solution in the presence of reduced CytC. The oxidation of reduced CytC by CIV was assayed via spectrophotometric measurement of reduced CytC at 550nm over 15 minutes. The reaction endpoint was assessed by artificially oxidizing all reduced CytC in the reaction mixture via the addition of the potent oxidizer potassium ferricyanide (K_3_Fe(CN)_6_), and the total reduced CtyC in the reaction was quantified using a triplicate of blank wells lacking tissue homogenate.
- Protein Concentration: Protein concentration of tissue homogenates was assayed using the Pierce BCA Protein Assay (Millipore).

References:

1.	Lanza, I.R., et al., *Chronic caloric restriction preserves mitochondrial function in senescence without increasing mitochondrial biogenesis.* Cell Metab, 2012. **16**(6): p. 777-88.

2.	Gnaiger, E., *Capacity of oxidative phosphorylation in human skeletal muscle: new perspectives of mitochondrial physiology.* Int J Biochem Cell Biol, 2009. **41**(10): p. 1837-45.

3.	Anderson, E.J., et al., *Mitochondrial H2O2 emission and cellular redox state link excess fat intake to insulin resistance in both rodents and humans.* J Clin Invest, 2009. **119**(3): p. 573-81.

**Supplementary table 1:** Linear regression models adjusting for age

|  |  | Coefficient Estimate [95% CI] | P value |
| --- | --- | --- | --- |
| *Model 1* JO_2_ (pmol/s/mg tissue)-State 3 (CI+II) | Intercept | 25.94 [17.62, 34.26] | <0.0001*** |
|  | IBM | -6.08 [-10.65, -1.50] | 0.01* |
|  | Age | -0.01 [-0.17, 0.15] | 0.89 |
| *Model 2* JO_2_ (pmol/s/mg tissue)-State 3 (CII) | Intercept | 16.90 [11.75, 22.05] | <0.0001*** |
|  | IBM | -3.69 [-6.53, -0.86] | 0.01* |
|  | Age | -0.01 [-0.11, 0.08] | 0.82 |
| *Model 3* JO_2_ (pmol/s/mg tissue)-Uncoupled Respiration | Intercept | 16.63 [11.11, 22.15] | <0.0001*** |
|  | IBM | -3.57 [-6.55, -0.58] | 0.02* |
|  | Age | -0.01 [-0.11, 0.09] | 0.84 |
| *Model 4* RCR | Intercept | 7.75 [6.21, 9.28] | <0.0001*** |
|  | IBM | -1.40 [-2.24, -0.56] | 0.001** |
|  | Age | -0.01 [-0.04, 0.01] | 0.19 |
| *Model 5* Leak Control Ratio | Intercept | 0.19 [0.11, 0.27] | <0.0001*** |
|  | IBM | 0.06 [0.02, 0.11] | 0.002** |
|  | Age | 0.0008 [-0.0007, 0.0024] | 0.28 |
| *Model 6* Maximal H_2_O_2_ emission  (%) | Intercept | 2.81 [1.32, 4.30] | <0.001*** |
|  | IBM | -1.47 [-2.29, -0.65] | <0.001*** |
|  | Age | 0.0092 [-0.02, 0.03] | 0.53 |
| *Model 7* TAC  (μM CRE / μg protein) | Intercept | 14.55 [-16.35, 45.47] | 0.34 |
|  | IBM | 15.27 [-1.15, 31.71] | 0.06 |
|  | Age | 0.13 [-0.45, 0.73] | 0.64 |
| *Model 8* CS activity (mU/mg of tissue) | Intercept | 2.91 [0.77, 5.04] | 0.0082** |
|  | IBM | -1.34 [-2.28, -0.40] | 0.006** |
|  | Age | 0.05 [0.01, 0.09] | 0.0092** |
| *Model 9*  mtDNA content | Intercept | 2331.33 [1521.05, 3081.61] | <0.001*** |
|  | IBM | -1395.15 [-1743.63, -1064.67] | <0.001*** |
|  | Age | 4.07 [-9.59, 17.73] | 0.55 |
| *Model 10* p-S65-Ub | Intercept | 20.64 [-107.32, 148.61] | 0.74 |
|  | IBM | 112.01 [67.82, 156.20] | <0.001*** |
|  | Age | 0.92 [-1.19, 3.05] | 0.38 |
| Abbreviations: CI, confidence interval; IBM, inclusion body myositis; JO_2_, oxygen flux; RCR, respiratory control ratio; TAC, total antioxidant capacity; CS, citrate synthase; p-S65-Ub, phosphorylated ubiquitin (S65). | | | |

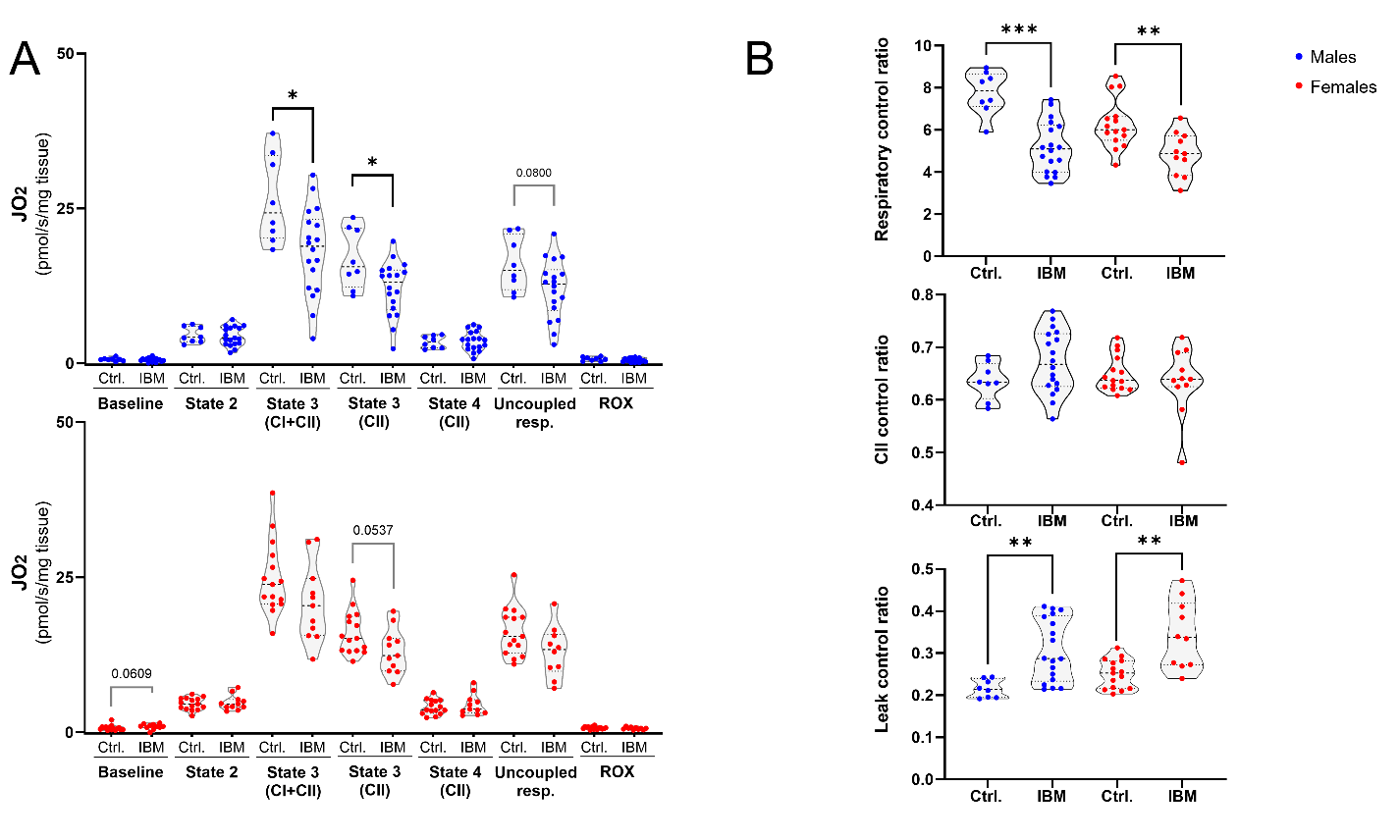

**Supplemental Figure 1: High resolution respiration in permeabilized muscle fibers.** A) Results segregated by sex for oxygen consumption rates normalized by tissue weight at each respiratory state. Similar patterns are seen in both males and females, although findings in females do not reach statistical significance, probably due to sample size. B) Flux control ratios segregated by sex.

*Violin plots show individual data with dashed lines representing medians and interquartile ranges. Group comparisons done using the Mann-Whitney tests. *P < 0.05, **P < 0.01, ***P < 0.001 ****P < 0.0001. Abbreviations: CI, Complex I; CII, Complex II; Ctrl, controls; J_O2_, oxygen consumption rate; IBM, inclusion body myositis; ROX, residual oxygen consumption*

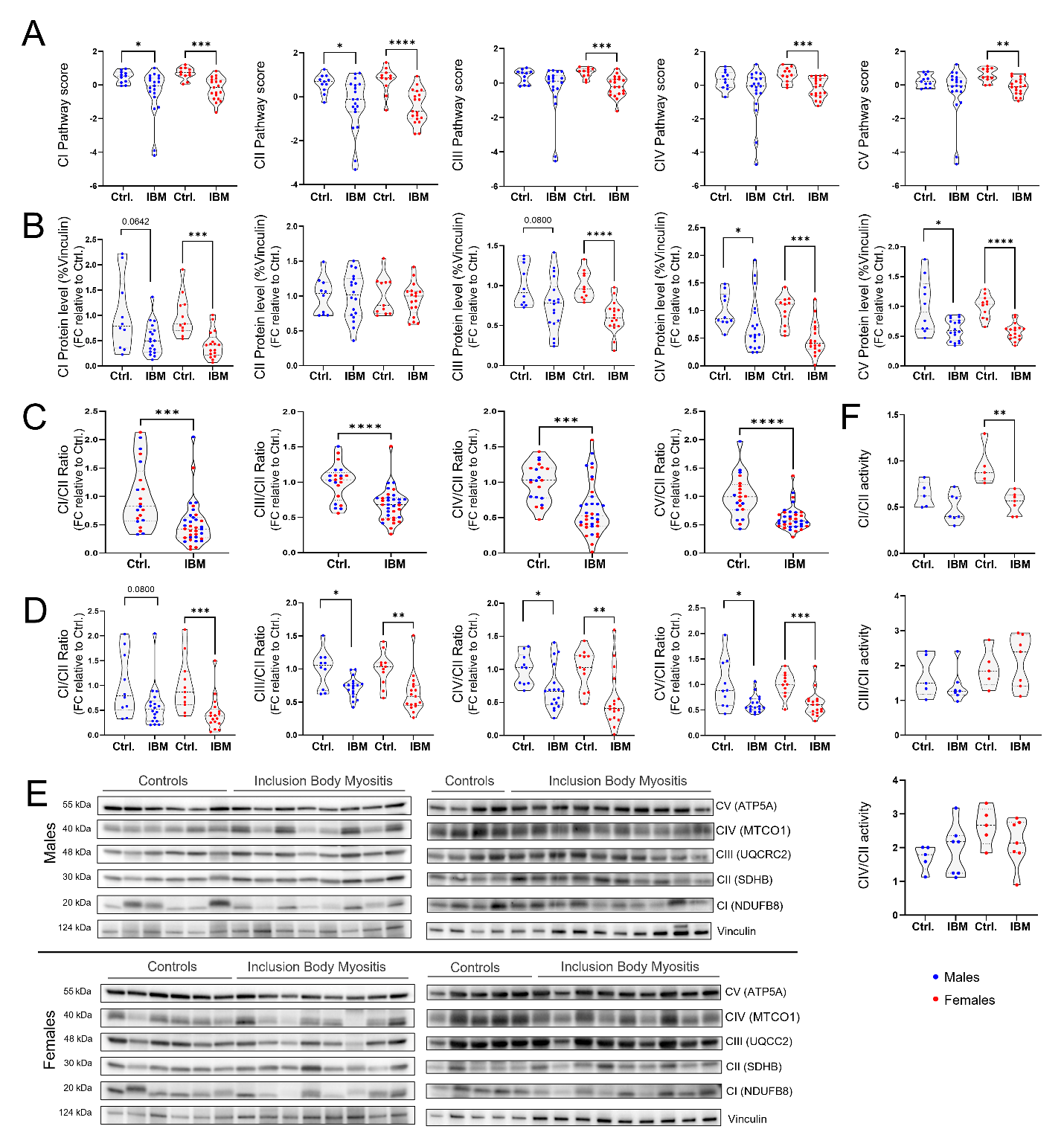

**Supplemental Figure 2: RNA expression, protein abundance and activity of OXPHOS complexes** A) Respiratory chain complex pathway scores by sex, B) fold change protein abundance (%Vinculin) by sex. C) Complex protein abundance relative to Complex II level, represented as fold change from control means, and D) segregated by sex. E) Western blot membranes of OXPHOS proteins on the retrospective (Left) and prospective (Right) cohorts. (F) Complex activity relative to CII activity segregated by sex.

*Violin plots show individual data with dashed lines representing medians and interquartile ranges. Group comparisons were done using the Mann-Whitney tests. *P < 0.05, **P < 0.01, ***P < 0.001 ****P < 0.0001, +/-. P values between 0.05 and 0.1. CI, Complex I; CII, Complex II; CIII, Complex III; CIV, Complex IV; CV, Complex V; Ctrl, controls; FC, fold change; IBM, inclusion body myositis.*

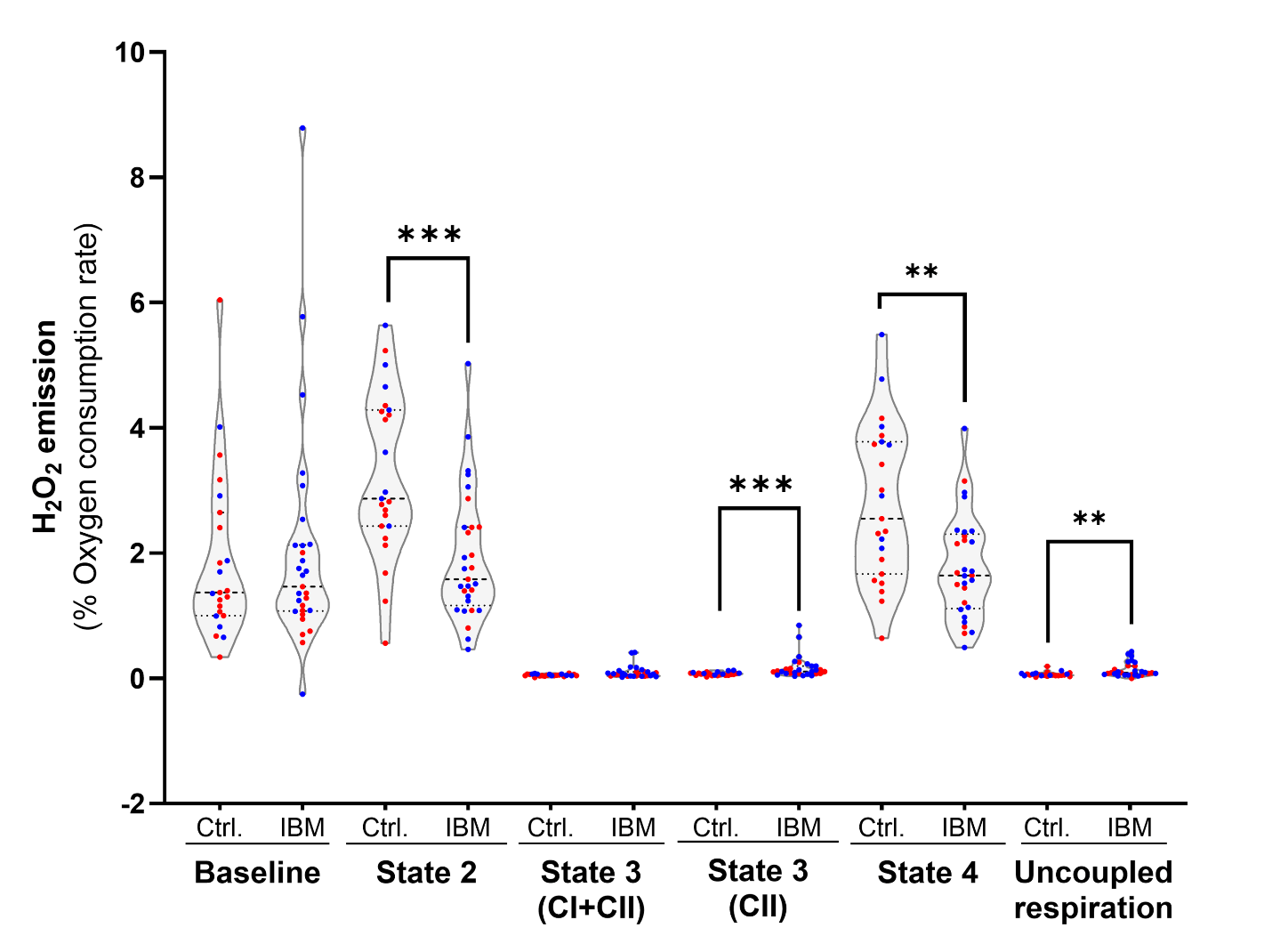

**Supplemental Figure 3:** **Reactive oxygen species production in IBM**. Detailed results of H_2_O_2_ emission during all respiratory states measured during Oroboros respirometry, normalized to oxygen consumption at each state.

*Violin plots show individual data with dashed lines representing medians and interquartile ranges. Group comparisons done using the Mann-Whitney tests. **P < 0.01, ***P < 0.001. Abbreviations: CI, Complex I; CII, Complex II; Ctrl, controls; IBM, inclusion body myositis*

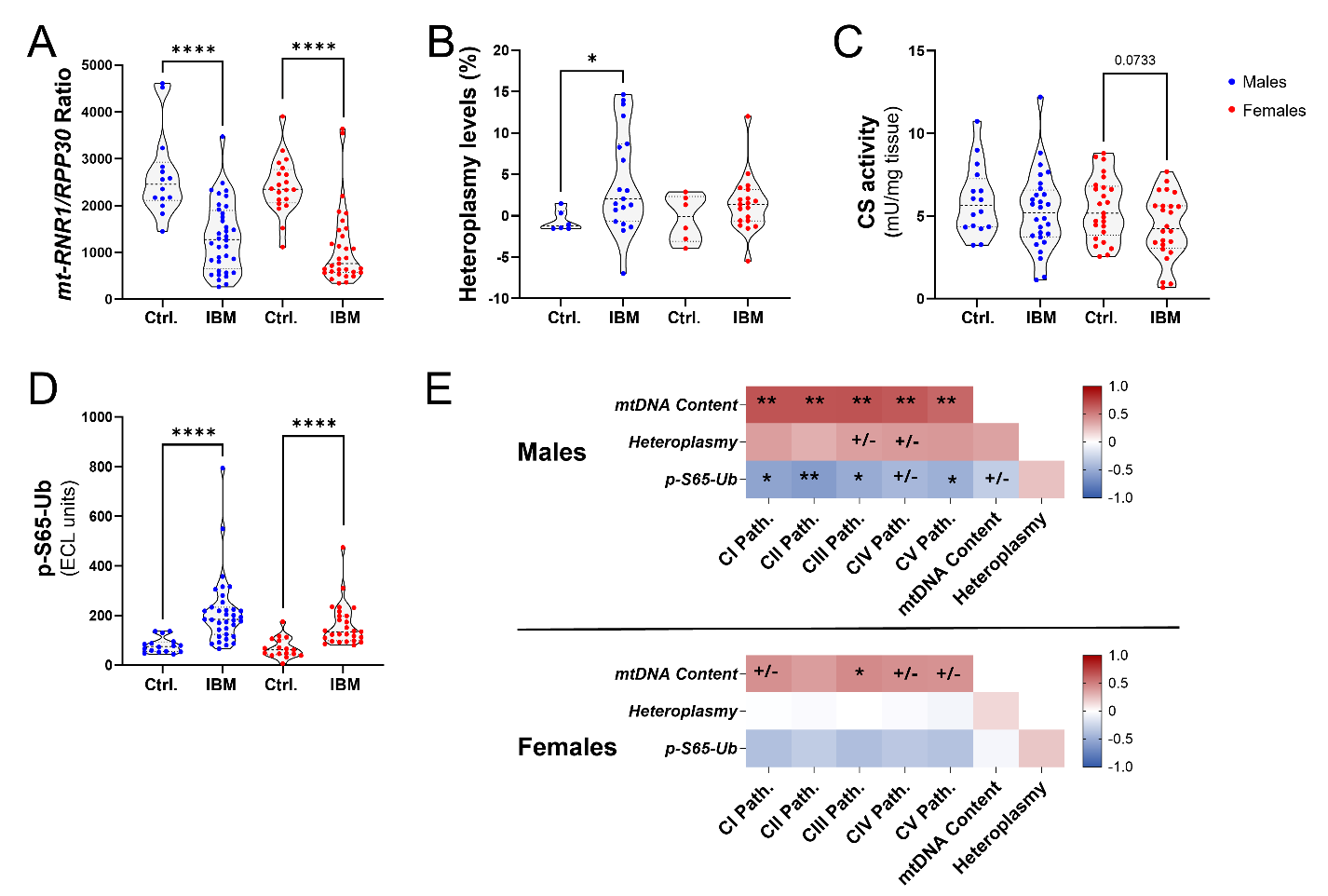

**Supplemental Figure 4**: **Mitochondrial DNA abnormalities, and mitophagy defects in inclusion body myositis** A) Mitochondrial DNA content (*mt-RNR1/*RPP30), B) Mitochondrial DNA deletion heteroplasmy levels C) Citrate synthase activity, and D) p-S65-Ub levels segregated by sex. E) Spearman correlations between respiratory complex RNA pathway scores, mtDNA content, mtDNA deletion heteroplasmy levels and p-S65-Ub levels in males (Top) and females (Bottom).

*Violin plots show individual data with dashed lines representing medians and interquartile ranges. Group comparisons were done using the Mann-Whitney tests. *P < 0.05, **P < 0.01, ****P < 0.0001, +/-. P values between 0.05 and 0.1. Abbreviations: CI, Complex I; CII, Complex II; CS, citrate synthase activity; Ctrl, controls; J_O2_, oxygen consumption rate; mtDNA, mitochondrial DNA; path., pathway score; RCR, respiratory control ratio*
